# Mechanical regulation of retinal vascular inflammation and degeneration in diabetic retinopathy

**DOI:** 10.1101/2022.12.18.520943

**Authors:** Sathishkumar Chandrakumar, Irene Santiago Tierno, Mahesh Agarwal, Emma M. Lessieur, Yunpeng Du, Jie Tang, Jianying Kiser, Xiao Yang, Timothy S. Kern, Kaustabh Ghosh

**Author notes:** Address all correspondence to: Kaustabh Ghosh, Ph.D. Department of Ophthalmology, University of California, Los Angeles, Doheny Eye Institute, 150 N. Orange Grove Blvd. Pasadena, CA 91103.

## Abstract

Vascular inflammation is known to cause degeneration of retinal vessels in early diabetic retinopathy (DR). Past studies investigating these diabetes-induced vascular abnormalities have focused primarily on the role of molecular or biochemical cues. Here we show that retinal vascular inflammation and degeneration in DR are also mechanically regulated by retinal vascular stiffening that is caused by overexpression of collagen-crosslinking enzyme lysyl oxidase (LOX) in retinal vessels. Treatment of diabetic mice with LOX inhibitor BAPN prevented the increase in retinal vascular stiffness, vascular ICAM-1 overexpression, and leukostasis. Consistent with these anti-inflammatory effects, BAPN treatment of diabetic mice blocked the upregulation of proapoptotic caspase-3 in retinal vessels, which concomitantly reduced retinal vascular degeneration and the diabetes-induced loss of contrast sensitivity in these mice. Finally, we show that increasing substrate stiffness alone increases the adhesiveness and neutrophil elastase-induced death of cultured retinal endothelial cells. By uncovering a link between LOX-dependent vascular stiffening and the development of retinal vascular and functional defects in diabetes, these findings offer unique insights into DR pathogenesis that has important translational potential.

## INTRODUCTION

Retinal vascular degeneration and loss of contrast sensitivity are clinical hallmarks of the early stages of diabetic retinopathy (DR). Retinal vascular inflammation, characterized by endothelial ICAM-1 upregulation and leukostasis, is strongly implicated in the development of these retinal abnormalities (1). Past work in DR has identified several molecular or biochemical factors as important mediators of these inflammatory changes. However, accumulating evidence from other inflammatory conditions such as sepsis and atherosclerosis indicate that vascular inflammation is also independently regulated by vascular stiffening caused by upregulation of collagen-crosslinking enzyme lysyl oxidase (LOX) (2, 3). This effect is attributed to endothelial mechanotransduction wherein alterations in cell or subendothelial matrix stiffness are transduced into intracellular biochemical signals that regulate endothelial cell (EC) function at the transcriptional and/or translational levels (4, 5). Since diabetes leads to increased LOX accumulation in the retina and vitreous of animal and human subjects, respectively (6, 7), here we asked whether diabetes-induced retinal vascular inflammation and associated retinal vascular and functional defects are mechanically regulated by LOX-dependent retinal vascular stiffening.

## RESULTS AND DISCUSSION

### LOX promotes retinal vascular stiffening and leukostasis in short-term diabetes

To assess the role of LOX in vascular stiffening-mediated retinal inflammation in diabetes, we first measured LOX expression in retinal vessels freshly isolated from nondiabetic and streptozotocin (STZ)-induced short-term (10 weeks duration) diabetic mice. RT-qPCR analysis revealed a 2.7-fold increase (*p*<0.0001; **(Fig.1A)** in retinal vascular LOX mRNA in diabetes. Importantly, this diabetes-induced LOX upregulation correlated with a significant increase (by 2.6-fold; p<0.0001; **Fig. 1B**) in retinal vascular stiffness, as measured by a biological-grade atomic force microscope (AFM). That LOX plays a crucial role in this retinal vascular stiffening was confirmed when treatment of diabetic mice with a specific and irreversible LOX inhibitor β-aminopropionitrile (BAPN) concomitantly and significantly reduced retinal vascular LOX mRNA (by 75%; p<0.001; **(Fig. 1A)** and vascular stiffness (by 90%; p<0.0001; **Fig. 1B**).

**Figure 1:**
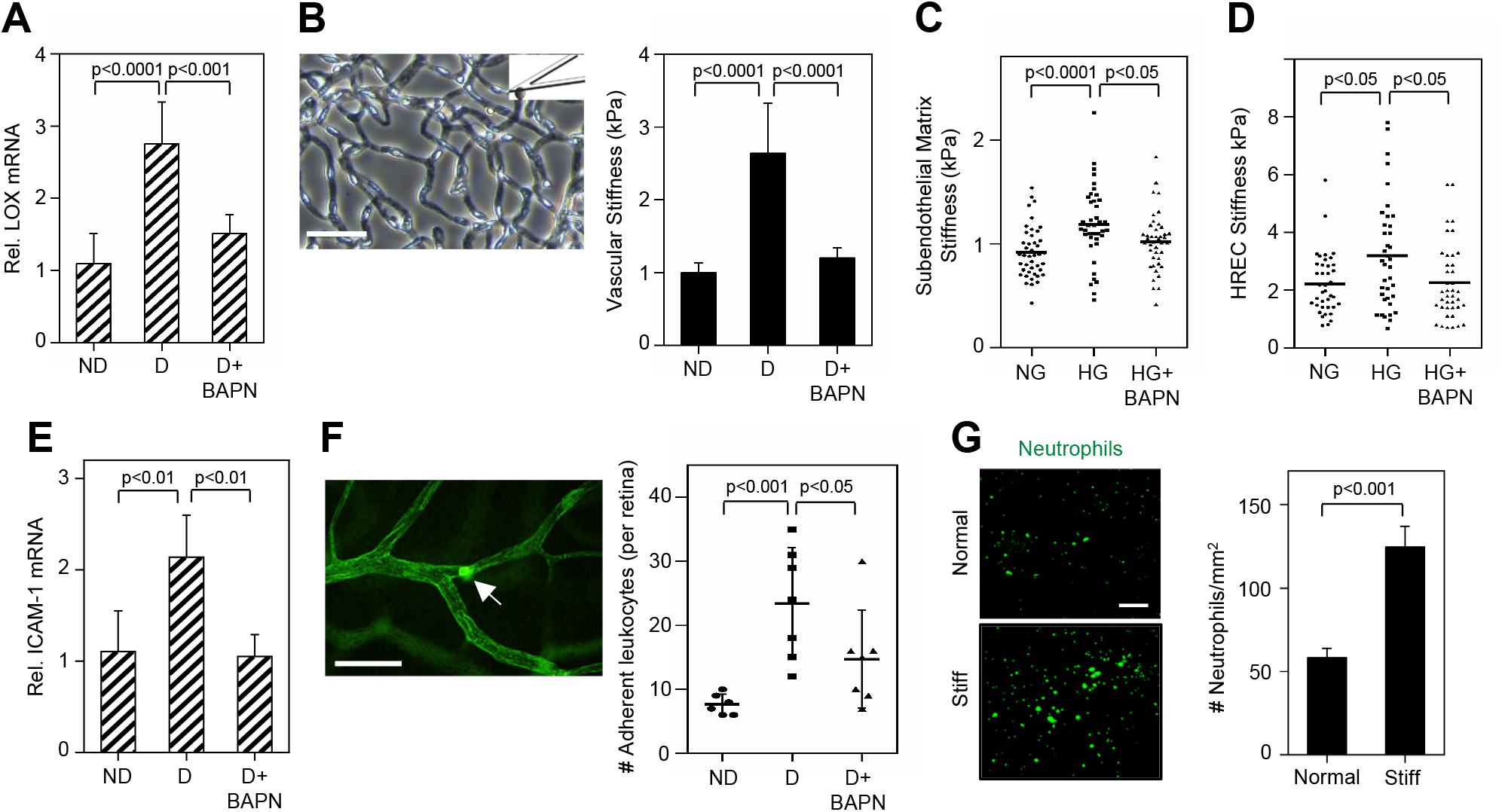
LOX promotes retinal vascular stiffening and leukostasis in short-term diabetes. **(A)** RT-qPCR analysis shows LOX mRNA levels (normalized w.r.t GAPDH) in freshly isolated mouse retinal vessels from nondiabetic (ND), diabetic (D), and BAPN-treated diabetic (D+BAPN) groups. **(B)** Stiffness of isolated mouse retinal vessels (shown in a representative image) was measured using a biological grade AFM fitted with a 1 um diameter probe. Data were obtained from ≥30 locations on the vasculature. Scale bar: 50 μm. **(C, D)** Stiffness of unfixed decellularized subendothelial matrix (C) or cultured HRECs (D) was measured using an AFM probe of 70 nm or 1 um diameter, respectively. Data were obtained from multiple locations on the matrix (n≥40) or multiple cells (n≥20). **(E)** RT-qPCR analysis shows ICAM-1 mRNA levels (normalized w.r.t GAPDH) from freshly isolated mouse retinal vessels. **(F)** Leukocyte adhesion to mouse retinal vessels (leukostasis) was assessed by injecting fluorescein-coupled concanavalin A lectin, followed by perfusion, whole-mount fluorescence imaging, and quantification of adherent cells (from 6-8 mice/group). Representative image shows a fluorescently-labeled adherent leukocyte (arrow). Scale bar: 50 μm. **(G)** Representative fluorescent images show dHL-60 neutrophils bound to HREC monolayers grown on normal (1 kPa) or stiff (2.5 kPa) synthetic matrices. Cell counts were obtained using Image J from ≥6 images/condition. Scale bar: 200 μm. Bars indicate mean ± SD for *in vivo* data (from 6-8 mice/group) or mean ± SEM for *in vitro* data.

Since LOX crosslinks collagen, we reasoned that the LOX-dependent retinal vascular stiffening in diabetes results from an increase in the crosslinking/stiffness of collagen-rich subendothelial matrix (basement membrane). Indeed, AFM measurement of decellularized subendothelial matrix derived from human retinal EC (HREC) culture demonstrated that LOX contributes greatly to the high glucose (HG)-induced matrix stiffening because treatment with LOX inhibitor BAPN reduced matrix stiffening by 60% (p<0.05) **(Fig. 1C)**. Further, consistent with the principle of mechanical reciprocity where adherent cells sense and respond to changes in matrix stiffness by undergoing a similar change in their own stiffness (8), we found that inhibition of subendothelial matrix stiffening by BAPN simultaneously prevents the HG-induced increase in HREC stiffness **(Fig. 1D)**. Thus, we propose that LOX increases retinal vascular stiffness in diabetes by crosslinking/stiffening the basement membrane (subendothelial matrix) that, via reciprocal interaction, simultaneously causes stiffening of the overlying retinal ECs.

To determine the inflammatory effects of LOX-dependent retinal vascular stiffening in diabetes, we measured ICAM-1 mRNA expression in freshly isolated retinal vessels. ICAM-1 is a crucial leukocyte adhesion molecule whose upregulation in retinal vascular ECs enhances leukocyte-dependent retinal vascular degeneration in early DR (9). As shown in **Fig. 1E–F**, the 2-fold increase (p<0.01) in retinal vascular ICAM-1 mRNA in short-term diabetic mice was completely blocked by LOX inhibition, which predictably reduced leukocyte adhesion to retinal vessels (leukostasis).

Notably, LOX exists in both soluble and matrix-localized forms, with the latter form implicated in matrix crosslinking and tissue stiffening (10). To determine whether matrix LOX-mediated vascular stiffening is sufficient to promote leukostasis independent of any potential inflammatory effects of soluble LOX, we assessed neutrophil adhesion to HRECs plated on ‘synthetic’ matrices of tunable stiffness that were fabricated to mimic retinal vascular stiffness of nondiabetic (1 kPa) or diabetic (2.5 kPa) mice (*refer to Fig. 1B*). As shown in **Fig. 1G**, HRECs plated on the stiff (2.5 kPa) matrix exhibited a 2-fold increase (*p*<0.001) in neutrophil adhesion than those grown on normal (1 kPa) matrix.

Collectively, these findings establish that diabetes-induced LOX upregulation in retinal vessels leads to vascular stiffening that, in turn, promotes vascular inflammation by increasing ICAM-1-dependent leukocyte adhesion to retinal ECs.

### LOX-dependent vascular stiffening enhances retinal endothelial apoptosis and vascular degeneration in long-term diabetes

Leukostasis is strongly implicated in retinal EC apoptosis that gradually leads to retinal vascular degeneration, a major clinically recognized lesion of early DR (11). Since we found that LOX-dependent vascular stiffening promotes retinal leukostasis in short-term diabetic mice, we asked whether LOX could thereby mechanically induce retinal EC apoptosis and vascular degeneration following prolonged diabetes (20-30 weeks duration). To address this question, we first confirmed that the retinal vessels of long-term diabetic mice continue to exhibit significant stiffening, which can be blocked by LOX inhibition **(Fig. 2A)**. Importantly, the stiffer retinal vessels in diabetic mice exhibited a 1.7-fold increase (p<0.05) in the mRNA levels of pro-apoptotic marker caspase-3, which was blocked by LOX inhibition **(Fig. 2B)**. Crucially, this BAPN-mediated suppression of caspase-3 mRNA was associated with a concomitant inhibition (by 50%; p<0.05) of the diabetes-induced increase in retinal vascular degeneration **(Fig. 2C)**.

**Fig.2:**
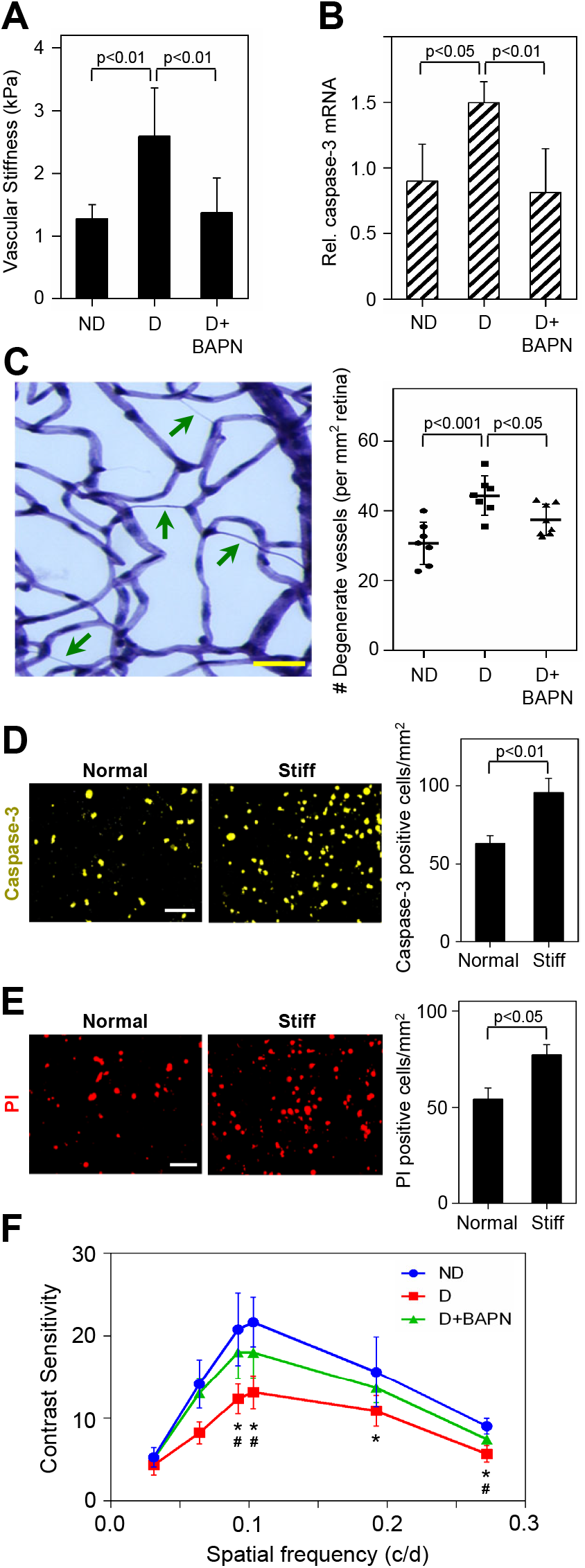
LOX-dependent vascular stiffening enhances retinal endothelial apoptosis and vascular degeneration in long-term diabetes. **(A)** Stiffness of isolated mouse retinal vessels was measured using a biological grade AFM fitted with a 1 um diameter probe. Data were obtained from ≥30 locations on the vasculature. **(B)** RT-qPCR analysis shows caspase-3 mRNA levels (normalized w.r.t GAPDH) from freshly isolated mouse retinal vessels. **(C)** Representative image shows degenerate retinal vessels (arrows) in a diabetic mouse. Number of degenerate vessels (from 6-8 mice/group) were counted and normalized w.r.t vascular area. Scale bar: 100 μm. **(D, E)** Representative fluorescent images show HRECs grown on normal or stiff synthetic matrices that were treated with neutrophil elastase for 12 h and labeled with caspase-3 activity reporter dye (D) or propidium iodide (PI) (E) to identify apoptotic or lysed cells, respectively. Counts of caspase-3- and PI-positive cells were normalized w.r.t image area. Bars indicate mean ± SEM for *in vitro* data and mean ± SD for *in vivo* data (from 6-8 mice/group). **(F)** Contrast sensitivity was measured at the same spatial frequencies (0.031 to 0.272 cycles/degrees) in all groups of mice following long-term diabetes. Line graphs indicate mean ± SD from 6-8 mice/group. *indicates p<0.001 for D vs ND; ^#^indicates p<0.01 for D vs D+BAPN.

Alteration in vascular/EC stiffness regulates EC fate by modifying its susceptibility to soluble factors (2, 12). Thus, we asked whether increased retinal vascular/EC stiffness in diabetes promotes EC apoptosis by increasing cell susceptibility to neutrophil elastase, a serine protease that is strongly implicated in neutrophil-induced retinal EC apoptosis (11). Indeed, neutrophil elastase treatment of HRECs grown on the stiff matrix, which mimics retinal vascular stiffness in diabetic mice, led to a 1.5-fold increase (with reference to/w.r.t normal matrix; p<0.01) in caspase-3 activation **(Fig. 2D)** that was predictably associated with a concomitant 40% increase (p<0.05) in propidium iodide (PI) labeling of dead HRECs **(Fig. 2E)**. Collectively, these findings establish that LOX-dependent retinal vascular stiffening contributes greatly to leukocyte-induced retinal EC death and vascular degeneration associated with early DR.

Loss of contrast sensitivity is a hallmark of early diabetes that correlates with DR severity (13). Mice exhibit this retinal defect within two months of diabetes, which exacerbates up to 10 months (14). Since the loss of contrast sensitivity in diabetic mice is expected to occur concomitantly with our observed LOX-mediated retinal inflammation and vascular degeneration, we asked whether LOX upregulation could also impair this retinal function in diabetes. Our findings revealed that diabetes predictably caused a substantial loss of contrast sensitivity in almost all spatial frequencies while, remarkably, LOX inhibition significantly prevented this vision defect **(Fig. 2F)**. Although this finding further underscores a crucial role for LOX in diabetes-induced retinal abnormalities, future studies will be needed to determine whether LOX impairs retinal function directly via a neuron-specific effect or indirectly via its mechanical regulation of retinal vascular inflammation, or both.

In summary, these findings have introduced a new paradigm for DR pathogenesis that identifies LOX-dependent vascular stiffening as a crucial and independent regulator of retinal vascular inflammation and degeneration in diabetes. A deeper understanding of the mechanical regulation of DR has the potential to identify entirely new classes of anti-inflammatory targets for more effective DR therapies in the future.

## MATERIALS AND METHODS

Detailed description of the methods is provided as Supporting Information.

### Experimental Animals

Diabetes was induced in adult C57BL/6J mice by streptozotocin injection using our previously reported protocol (6). LOX inhibitor β-Aminopropionitrile (BAPN) was administered in diabetic mice through drinking water. Mice were euthanized at 10 weeks (short term) or 20-30 weeks (long term) duration of diabetes and eyes were harvested for further analysis.

### Retinal Vascular Stiffness

Retinal vessels were isolated from mildly-fixed eyes following trypsin digestion and subjected to stiffness measurement with a NanoWizard^®^ 4 XP BioScience atomic force microscope (AFM; Bruker, Inc.) fitted with 1 μm hemisphere nitride probe. AFM Force Curves were obtained in the contact mode force spectroscopy mode and analyzed using JPK Data Processing Software.

### Statistics

All data were analyzed using GraphPad Prism 6.01 (GraphPad Software, San Diego, CA). Statistical differences between three or more groups were assessed using a one-way analysis of variance (ANOVA), followed by Tukey’s *post hoc* multiple comparisons test. Two-tailed unpaired Student’s *t* test was used for studies comparing two experimental groups. *p*<0.05 considered as statistically significant.

## ACKNOWLEDGMENTS

This work was supported by NIH grants R01EY028242 (to K.G.), R01EY033002, R01EY022938, and R24EY024864 (to T.S.K.), EY022938-S1 (to E.M.L.), and P30EY034070-01 (NEI core grant to the C.T.V.R at UC Irvine), The Stephen Ryan Initiative for Macular Research (RIMR) Special Grant from W.M. Keck Foundation (to Doheny Eye Institute), and Ursula Mandel Fellowship and UCLA Graduate Council Diversity Fellowship (to I.S.T.). This work was also supported by Unrestricted Grants from Research to Prevent Blindness, Inc. to the Department of Ophthalmology at UCLA and Gavin Herbert Eye Institute at UC Irvine.

