## Supporting Information for "Mechanical regulation of retinal vascular inflammation and degeneration in diabetic retinopathy"

### Extended material and methods

#### Experimental Animals

All animal studies were conducted in accordance with the Association for Research in Vision and Ophthalmology (ARVO) Statement for the Use of Animals in Ophthalmic and Vision Research and approved by the Institutional Animal and Care Use Committee (IACUC) at the University of California Riverside, University of California Irvine, and Case Western Reserve University. Diabetes was induced in adult (8 week-old) male C57BL/6J mice (Jackson Laboratory, Bar Harbor, ME, USA) through daily i.p. injections of freshly-prepared streptozotocin solution (STZ, MP Biomedicals, Irvine, CA; 60 mg/kg body weight in 10 mM citrate buffer; pH 4.5) for five consecutive days. Mice with fasting blood glucose >275 mg/dL two weeks after the last STZ injection were classified as diabetic, with this time point being considered as the onset of overt diabetes. Age-matched normal C57bL/6J mice receiving citrate buffer alone were classified as nondiabetic. LOX inhibition was achieved in diabetic mice by administering a specific and irreversible LOX inhibitor β-aminopropionitrile (1) (BAPN; Catalog no. A3134-5G; Sigma-Aldrich, St. Louis, MO, USA) through drinking water (3 mg/kg) for 10 weeks (short-term) or 20-30 weeks (long-term) prior to euthanasia and collection of eyes for subsequent analyses.

#### Isolation of Retinal Vessels

For gene expression analysis, retinal vessels were isolated from fresh (unfixed) eyes using the hypotonic method (2). Briefly, freshly-isolated whole retinas were incubated in de-ionized water for ~2h until they became fuzzy, then transferred (using an inverted glass Pasteur pipette) into a well containing 80 Kunitz units DNase (Catalog no. 79254; Qiagen, Hilden, Germany) for 5-10 min at 37°C. Neuronal debris was removed by repeated injection of DNase solution on the submerged retina to cause gentle agitation, resulting in isolation of clean retinal vessels. Two diaphanous retinal vasculatures were pooled and used for RT-qPCR analysis.

For stiffness measurements, retinal vessels were isolated using trypsin digestion method (3, 4). Briefly, enucleated eyes were mildly fixed in 5% formalin for 24 h at 4ºC prior to isolation of whole retinas, which were then subjected to 10% Trypsin digestion (Trypsin 1:250, Ameresco/VWR Life Science, PA) for up to 3h. The isolated vessels where then transferred to and rinsed in $ddH_2O$ followed by mounting on cleaned Superfrost™ Plus microscope slide (Fisher) for stiffness measurement.

**RT-qPCR**

Total RNA was isolated from retinal vessels using Direct-zol™ RNA MiniPrep (Zymo Research, Irvine, CA, USA) and converted to cDNA using High-Capacity RNA-to-cDNA kit (catalog no. 4387406; Thermo Fisher Scientific-Applied Biosystems™) prior to amplification with gene specific Taqman primers for LOX (Mm00495386_m1), ICAM-1 (Mm00516023_m1), and Caspase-3 (Mm01195085_m1) in QuantStudio™ 5 Real-Time PCR system. The target gene expression was normalized to the house keeping gene GAPDH (Mm99999915_g1) and relative expression was determined using comparative Delta Delta Ct (DDCt) method. Negative control was performed for each reaction assay plate.

**Cell culture and treatments**

Human retinal endothelial cells (HRECs) were purchased (catalog no. ACBRI 181; Cell Systems corp., Kirkland, WA, USA) and cultured as described previously (5). Human promyelocytic leukemia cell line (HL-60) was purchased from ATCC (Manassas, VA, USA) and cultured in RPMI 1640 medium (Gibco™-Thermo Fisher Scientific) supplemented with 10% fetal bovine serum (FBS; Catalog no. SH30396.03, Hyclone, Logan, UT) and Penicillin-Streptomycin (Gibco™-Thermo Fisher Scientific). HL-60 cells were differentiated with 1.4% DMSO for 5 days to acquire the functional properties of neutrophil (dHL-60) (6, 7) and used as an alternative to primary neutrophils (8) in neutrophil-HREC adhesion assay. ***In vitro treatment regimen***: Confluent HRECs (passages 5-8) were grown in regular culture medium containing either 5.5 mM glucose (normal glucose, NG) or high glucose (30 mM; HG) ± 0.1 mM BAPN for 15 days prior to measurement of cell stiffness or decellularization to obtain subendothelial matrix. These cultures were also supplemented with 200 µg/mL L-ascorbic acid (Catalog no. A4034; Sigma-Aldrich) to facilitate matrix deposition.

**Subendothelial matrix**

Decellularized subendothelial matrix was obtained as per our published protocol (9). Briefly, HRECs in NG or HG±BAPN medium were grown on activated glass coverslips for

15 days prior to decellularization with mild detergent that removes the cells while preserving the HREC-secreted matrix. Residual cell debris was removed from the decellularized matrix by DNase treatment prior to stiffness measurement.

**Stiffness measurement**

A NanoWizard® 4 XP BioScience atomic force microscope (AFM; Bruker Inc., Santa Barbara, CA), coupled with a Zeiss Axiovert phase contrast microscope, was used for stiffness measurement of retinal vessels, subendothelial matrix, and HREC cultures. For measurement of retinal vascular or HREC stiffness, AFM was fitted with a pre-calibrated SSA-SPH-1UM probe (spring constant 0.25 N/m; Bruker AFM Probes, CA, USA) containing a 1 μm-radius hemispherical silicon nitride tip. For measurement of subendothelial matrix stiffness, a pre-calibrated PFQNM-LC-A-CAL probe (spring constant 0.075 N/m; Bruker AFM Probes, CA, USA) containing a 70 nm-radius hemispherical silicon nitride tip was used. Stiffness was measured in the contact mode force spectroscopy mode by applying a 50-200 pN (set point) indentation force. Multiple force curves ($n \geq 30$/group for retinal vessels; $n \geq 20$/group for HRECs; $n \geq 40$/group for matrix) were analyzed using JPK Data Processing Software.

**Leukostasis**

Retinal leukostasis (leukocyte adhesion to retinal vessels) was investigated in nondiabetic and short-term (10 weeks duration) diabetic ± BAPN mice, as previously reported (10). Briefly, mice were anesthetized using i.p. injection of a solution containing 17.5 mg/mL ketamine and 2.5 mg/mL xylazine and exsanguinated by perfusion with 0.9% (w/v) sodium chloride (Valley Vet Supply, Marysville, KS, USA) for 2 min. Fluorescein-coupled concanavalin A lectin (cat. no. FL-1001-25, Vector Laboratories, Burlingame, CA, USA) at 20 μg/ml in 0.9% sodium chloride was then infused as described previously (10). Eyes were enucleated and flat mounted retinas were imaged using Nikon Eclipse Ni wide-field epifluorescence microscope using a green filter (395/509 nm). Brightly fluorescent leukocytes were counted and plotted as total number of adherent leukocytes per retina.

**Synthetic matrix fabrication**

Synthetic matrices of tunable stiffness were fabricated using polyacrylamide, as previously reported (11, 12). Briefly, thin (~100 µm thick) elastic synthetic matrices of 1 kilopascal (1 kPa; 'normal') and 2.5 kPa ('stiff') stiffness were fabricated to mimic retinal vascular stiffness in nondiabetic and diabetic conditions, respectively, prior to surface coating with the proinflammatory matrix molecule fibronectin (5 µg/cm$^2$). HRECs were grown on these synthetic matrices in regular culture medium containing 5.5 mM glucose for use in functional assays.

**Neutrophil-EC adhesion**

Neutrophil-EC adhesion assay was performed as per our previously reported protocol (13, 14). Briefly, HRECs were grown to confluence on synthetic matrices for 24h before addition of fluorescently-labeled neutrophils at 125,000 cells/cm$^2$ for 30 min at 37°C in serum starvation medium (2.5% fetal bovine serum). After washing away the non-adherent neutrophils with PBS, the co-culture was fixed with 1% PFA and the adherent neutrophils imaged using a Zeiss Axio Observer microscope and counted using ImageJ ($n \geq 6$ images per condition). Total number of adherent neutrophils were normalized w.r.t area (mm$^2$) and expressed as #neutrophils/condition.

**Caspase-3 activation and EC death**

To determine the role of matrix stiffness in retinal EC apoptosis and death associated with early DR, HRECs (25k/cm$^2$) were plated on polyacrylamide-based 'normal' (1 kPa) or 'stiff' (2.5 kPa) synthetic matrices for 48h in culture medium followed by treatment with human neutrophil elastase (50nM) for 12h in starvation medium (2.5% FBS). HREC apoptosis (caspase-3 activation) was monitored in real time by adding Biotium NucView 488 Caspase-3 substrate (Biotium; Fremont, CA, USA) to HRECs together with neutrophil elastase and imaging using the Zeiss Axio Observer microscope. At the end of 12h treatment, propidium iodide (Thermo Fisher Scientific) was added to the cultures to detect membrane lysis (cell death) prior to imaging. Caspase-3- and PI-positive cells were counted using ImageJ ($n \geq 6$ images per condition) and normalized w.r.t area (mm$^2$).

**Retinal vascular degeneration**

Retinal vascular degeneration was assessed in nondiabetic and longer-term (30 weeks duration) diabetic ± BAPN mice, as previously described (10). Briefly, eyes were removed and fixed in 10%

formalin for 10 days. Retinas were then isolated and digested in elastase (pH 6.5) for 2 h at 37°C followed by overnight incubation of the vasculature in 100 mM Tris buffer (pH 8.5) at room temperature. Non-vascular cells were gently brushed away under a dissecting microscope and the cleaned retinal vasculature was spread on a Superfrost™ Plus microscope slide and allowed to air dry overnight before staining with hematoxylin and periodic acid-Schiff reagent. Acellular vessels, identified as capillary-sized tubes having no nuclei anywhere along their length, were counted from approximately seven field areas around the mid-retina and plotted as number of degenerate vessels per area.

**Contrast Sensitivity**

Contrast sensitivity was measured in nondiabetic and longer-term (30 weeks duration) diabetic ± BAPN mice as previously reported (15, 16). Briefly, mice were subjected to a virtual optokinetic test where their ability to track the movements of grating lines of varying contrast was assessed at six different spatial frequencies viz. 0.031, 0.064, 0.092, 0.103, 0.192, and 0.272 cycles/degree (c/d). Contrast sensitivity, which reflects the functional response of spatially sensitive retinal cells, was determined as the inverse of Michelson contrast, without correction for luminance of the monitors. Animals were not anesthetized and placed in the optokinetic device for training prior to recording their response. Each measurement was repeated multiple times to assess the reproducibility of responses.
